# The role of CLV1, CLV2 and HPAT homologs in nitrogen-regulation of root development

**DOI:** 10.1101/2020.08.01.232546

**Authors:** Chenglei Wang, James B Reid, Eloise Foo

## Abstract

Plants use a variety of hormonal and peptide signals to control root development, including in adapting root development to cope with nutrient stress. Nitrogen (N) is a major limiting factor in plant growth and in response to N stress plants dramatically modulate root development, including in legumes influencing the formation of N-fixing nodules in response to external N supply. Recently, specific CLE peptides and/or receptors important for their perception, including CLV1 and CLV2, have been found to play important roles in root development in a limited number of species, including in some cases the response to N. In the legume *Medicago truncatula*, this response also appears to be influenced by RDN1, a member of the hydroxyproline O-arabinosyltransferase (HPAT) family which can modify specific CLE peptides. However, it not known if this signalling pathway plays a central role in root development across species, in particular root responses to N. In this study, we sought to systematically examine the role of homologues of these genes in root development of the legume pea (*Pisum sativum*. L) and non-legume tomato (*Solanum lycopersicum*) using a mutant based approach. This included a detailed examination of root development in response to N in these mutant series in tomato. We found no evidence for a role of these genes in pea seedling root development. Furthermore, the *CLV1-like FAB* gene did not influence tomato root development, including N response. In contrast, both *CLV2* and the *HPAT FIN* appear to positively influence root size in tomato but do not mediate root responses to N. These suggest a relatively species-specific role for these genes in root development, including N regulation of root architecture.

## Introduction

Flexible root morphology is critical to plant survival to enable plants to adapt and respond to different environment conditions. Nutrient levels are one of the main factors that shapes the root system architecture (López-Bucio et al. 2003). Nitrogen (N) is particularly important in shaping root development as it is the major limiting nutrient for plant growth (Liu et al. 2019). Depending on the form of N (nitrate or ammonium) as a general rule local N supply can stimulate lateral root development to enhance foraging, while severe nitrogen deprivation limits lateral root formation (reviewed by Motte et al. 2019). Interestingly very high nitrogen levels can also limit lateral root elongation and/or number, although this is species specific (e.g. Zang et al. 2020; Zhang and Forde 1998). Many legumes can form a second secondary root organ, the nodule, which house nitrogen-fixing rhizobial bacteria and the formation of these nodule organs is dramatically repressed by nitrogen supply (Goh et al. 2018). Indeed, in legumes there is a clear overlap in some of the genes and signals that regulate root development and nodulation in response to nitrogen and this dovetails with recent evidence that legume nodules are derived in part from a modified lateral root organ development program (Schiessl et al. 2019; Soyano et al. 2019).

Studies seeking to understand the molecular and genetic mechanisms driving root development in responses to nitrogen, including nodulation, have revealed key roles for two parallel peptide hormone signalling pathways. Firstly, via the CEP peptide pathway, local nitrogen starvation inhibits lateral root growth through the induction of the production of CEP peptides, which are then transported to the shoot and perceived in the shoot by leucine rich repeat receptor like kinases (LRR-RLK) (including CEPR1 and 2). The signalling perception in the shoot then trigger a downstream signal to the root to stimulate lateral root growth in areas exposed to high nitrogen (e.g. Bisseling and Scheres 2014; Tabata et al. 2014). Secondly, elements of the CLV-CLE peptide pathway have been shown to be involved in mediating nitrogen response of roots. In legumes, this involves elements of the negative feedback autoregulation of nodulation (AON) pathway (reviewed by Wang et al. 2018). In AON, CLE peptides are induced by rhizobia and/or high nitrogen and are perceived by LRR-RLK in *Medicago truncatula, Lotus japonicus*, soybean and pea (SUNN/HAR1/NARK/SYM29). Mutants disrupted in these receptors display excess or supernodulation even in the presence of high nitrogen. In the absence of rhizobia and hence nodule formation, there is also a subtle but consistent effect of the *nark, sunn* and *har1* mutations on root architecture, including increased number of lateral roots and in the case of *har1* and *sunn* shorter primary roots (Day et al. 1986; Goh et al. 2018; Lagunas et al. 2019; Schnabel et al. 2012; Wopereis et al. 2000). In *M.truncatula*, several reports suggest *SUNN* is important for nitrogen regulation of root architecture and shoot/root biomass partitioning, although the specific traits influenced did vary between studies (Goh et al. 2018; Jin et al. 2012; Lagunas et al. 2019). *MtRDN1* and its orthologue *LjPLENTY*, which encode hydroxyproline O-arabinosyltransferases (HPAT) that modify some CLE peptides important for AON, have also been reported to positively influence root length in *M.truncatula* and *L.japonicus* and play a role in the nitrogen regulation of root development in *M. truncatula* (Goh et al. 2018; Schnabel et al. 2011; Yoshida et al. 2010). *L.japonicus* mutants disrupted in another AON acting LRR-RLK, KLV, exhibit reduced tap and lateral root lengths and a reduced number of lateral roots (Oka-Kira et al. 2005) but the role of this gene and other LRR-receptors involved in AON (CLV2, CRN) in nitrogen regulation of root development of legumes has not yet been characterised.

Interestingly, elements of the AON pathway outlined above are highly homologous to elements of the CLAVATA-WUSCHEL negative feedback loop that perceive CLE peptides to control shoot apex cell proliferation (reviewed by Kitagawa and Jackson 2019). Mutants in elements of this pathway in Arabidopsis, rice, maize and tomato have clear influences on shoot, flower and/or fruit development. There is some evidence that some CLE peptides and receptors (CLV1, CLV2, CRN) also play roles in regulating root architecture in non-legumes, in particular in response to nutrient deprivation. Applications of CLV3-like peptides to Arabidopsis suppress root growth and roles for CLV2, CRN and CLV1 in the perception of specific peptides have been reported (Fiers et al. 2005; Hobe et al. 2003; Pallakies and Simon 2014; Stahl et al. 2009; Whitewoods et al. 2018). Indeed, application of *Physcomitrella* CLE peptides can induce similar responses in Arabidopsis roots (Whitewoods et al. 2018) and application of *At*CLE26 also inhibits some aspects of root growth in rice, *Brachypodium* and tomato (Czyzewicz and De Smet 2016; Czyzewicz et al. 2015). In rice application studies have shown the CLE peptide FCP2 influences root apical meristem maintenance but the precise receptors have not yet been investigated (Chu et al. 2013). The CLV1 receptor is essential for regulation of lateral root number in response in nitrogen deprivation in Arabidopsis and the authors identified several CLE peptides important for this response (Araya et al. 2014). In addition, terminal differentiation of the root apical meristem in Arabidopsis by low phosphate involves CLV2 (Gutierrez-Alanis et al. 2017). However, only subtle changes in the properties of root apical meristems have been observed in *clv1* and higher order mutants of Arabidopsis (Stahl et al. 2013; Whitewoods et al. 2018). Indeed, unlike studies outlined in legumes above, mutations in receptors of the CLAVATA-WUSCHEL pathway in Arabidopsis do no display gross changes in root size or root architecture in the absence of CLE peptide application and/or under nutrient sufficient conditions (e.g. Fiers et al. 2005; Gutierrez-Alanis et al. 2017; Miwa et al. 2008; Pallakies and Simon 2014).

In non-legumes, although the role of CLV1 in N regulation of root development has been tested in Arabidopsis, the role of other components of the CLV-CLE peptide signalling pathway have yet to be investigated and how conserved this is across other non-legume species is not known. In addition, in legumes the influence of CLV-CLE pathway on specific aspects of root development, including in response to N, appears to vary between studies. To examine the centrality of CLV-CLE pathway in root development of legumes and non-legumes we systematically characterised the root development of a model legume (pea) and non-legume (tomato) disrupted in receptors encoded by *CLV1-like* genes (*PsNARK/SlFAB*) and *CLV2*, and in hydroxyproline O-arabinosyltransferase genes (*PsRDN1/SlFIN*) (Krusell et al. 2002; Krusell et al. 2011; Schnabel et al. 2011; Xu et al. 2015). In tomato, we examined in detail the role of not just CLV1 but also CLV2 and HPAT in N regulation of root development. Comparison of the legume pea and non-legume tomato is informative given that nodulation has a dramatic effect on the nitrogen economy of legumes and therefore adaptive responses in root phenotypes to nitrogen may be different between legumes and non-legumes.

## Materials and Methods

### Plant material

The tomato wild type (*Solanum lycopersicum* cv. M82) and the mutants on this background, *Slfab* (*Slclv1*), *Slclv2-2, Slclv2-5, Slfin-n2326* and *Slfin-e4489* were described in Xu et al (2015) and were provided by Prof Zachary (Cold Spring Harbor Laboratory, USA). For pea studies, the wild type was cv. Frisson and the mutants were *Psnark* (P88, formerly *Pssym29*), *Psclv2* (P64, formerly *Pssym28*) and *Psrdn1* (P79, formerly *Psnod3*) on this background (Duc and Messager, 1989; Sagan and Duc, 1996).

### Mature plant growth system in pots

To exclude rhizobia and prevent nodulation, pea plants were grown for 14 days under sterile conditions with no additional nutrients as outlined in McAdam et al. (2017). Briefly, pea seed was surface sterilised, nicked and planted in sterile vermiculite in controlled cabinets (20 °C day/15 °C night, 18 hour photoperiod, 150umolms-1 at pot height). Plants were watered as needed with UV-sterilised water.

Studies of mature tomato plants were grown in pots under non-sterile conditions. Tomato seeds were germinated in potting mix and transplanted to 2L pot two weeks after sowing. The tomato pots contained a 1:1 mixture of vermiculite and gravel topped with vermiculite. For the grafting experiment, wedge grafts were performed in the hypocotyl three days after transplanting the rootstocks. After grafting, the plants were maintained in a humid environment under shade cloth until scions showed new growth. The plants were then gradually reintroduced to ambient conditions (approx. 5 - 7 days after grafting). Plants were grown in glasshouse conditions under an 18 h photoperiod and at a temperature generally ranging from 13 °C to 21°C. For experiments using tomato *clv2* mutant plants, the plants were grown in a controlled glasshouse at 25 °C day/20 °C night and an 18 hour photoperiod. Tomato plants were supplied with 75ml of modified Long Ashton Nutrient Solution (LANS) 2-3 times a week containing 5 mM KNO_3_ and 0.5 mM NaH_2_PO_4_.

### In vitro *plate growth system for tomato seedlings*

To examine tomato seedling development, tomato seeds were grown under semi-sterile conditions on large sterile petri plates (23 × 23cm; Thermo Scientific). Initial root phenotyping of the *fab, clv2* and *fin* mutants was conducted using half strength LANS containing 2.5 mM NaH_2_PO_4_ and 5.6 mM KNO_3_ (Hewitt 1966) solidified with 5g phytagel/L (Sigma, USA). In subsequent experiments that aimed to examine the tomato root morphology response under different N conditions, the tomato seedlings were grown in modified MGRL media with 1.75mM HPO_4_^-^ (from a mixture of Na_2_HPO_4_ and NaH_2_PO_4_) and various NO_3_^-^ (from KNO3 and Ca(NO_3_)_2_·4H_2_O) concentrations solidified with 5g phytagel/L as outlined in Araya et al. (2014). For both media types, the pH of the solution was adjusted to 5.8 with NaOH or HCl before phytagel was added and the media autoclaved. Approximately 275ml medium was added to each plate, which were then placed on an angle to enable a 7cm media-free area at the top of the plate for shoot growth).

Tomato seeds were surface sterilised with 70% ethanol for 3 mins with shaking, then rinsed thrice in sterile Milli Q water. Three seeds per plate were sown directly onto media (8 - 9cm down from the top of the plate), the plates were sealed with micropore paper tape (Nexcare) and the bottom half of plates was covered with black plastic to reduce the exposure of the root to light. The plates were placed on a slight slope in a controlled glasshouse under an 18 h photoperiod, with 25°C day/20°C night temperatures.

### Scoring of root morphology

For pea experiments, plants were harvested and tap root length, total secondary lateral root number, length of the top six secondary lateral roots and internode length from nodes 1-5 was recorded. Shoot and root tissue was separated and dried at 60°C.

For pot grown tomato plants the root and shoot tissues were harvested 8 weeks after transplantation and dried at 60°C. At asses root apical meristem size, 12 root tips per plant were collected from 5 week old pot grown tomato plants, stained with 1% acetocarmine, inspected under microscope and the ImageJ software was used to estimate root apical meristem length and area (Suppl Fig 2). For plate grown tomato seedlings the tap root length and shoot length (the length from the root shoot junction to the top of the apex) were measured and the total number of secondary lateral roots counted. In the experiments to characterise the root phenotype of mutants, the length of the longest lateral root was measured and the number of tertiary roots on the longest secondary root was recorded. For the experiments with various N treatments, the length of the tap root with red colouration due to anthocyanin build up was also measured. At the conclusion of the experiment, the root and shoot tissues were harvested, washed and dried at 60°C. Two N experiments were conducted with mutants, one examining *fab* and wild type (Fig 6) and an independent experiment examining *clv2-5, fin-n2326* and wild type. For clarity of presentation, data for *clv2-5* and *fin-n2326* mutant plants is presented in separate figures, meaning the wild type data is the same in both Fig 7 and 8.

### Phylogenetic analyses

The multiple sequence alignment was generated using the Muscle algorithm in MEGA7 (Edgar 2004). The phylogenetic tree was constructed using the Maximum Likelihood method within the MEGA7 software (Kumar et al. 2016). The percentage of trees in which the associated taxa clustered together is shown next to the branches. Initial tree(s) for the heuristic search were obtained automatically by applying Neighbor-Join and BioNJ algorithms to a matrix of pairwise distances estimated using a JTT model, and then selecting the topology with superior log likelihood value. A discrete Gamma distribution was used to model evolutionary rate differences among sites (5 categories (+G, parameter = 1.6783)). The tree is drawn to scale, with branch lengths measured in the number of substitutions per site.

### Statistical analyses

The data was analysed using SPSS software (vision 20, IBM). The normal distribution of data and the homogeneity of variances was analysed with the Shapiro-Wilk test (P<0.05) and homogeneity test (P<0.05), respectively. When both tests were not significant, the data were subjected to either one-way or two-way ANOVA followed by a Tukey’s post-hoc test to compare the means of different groups (if there were more than 2 groups). For the data that were either not normally distributed or did not have equal error variance, the data were log or square transformed and AONVA analysed on transformed data.

## Results

### Phylogentic relationships amongst pea and tomato gene famalies

To examine the realtioships amongst legume and non-legume members of gene famalies, alignments were performed with tomato and pea, and other relevant legume and non-legume genes (Suppl. Fig 1). Analysis of *CLV1*-like sequences revealed the tomato *SlFAB* (Solyc04g081590) gene in a clade with Arabidopsis *AtCLV1*, rice *OsFON1* and legume AON *CLV1*-like genes including pea *PsNARK*, Medicago *SUNN* and Lotus *HAR1* (Suppl. Fig 1b). Analysis of the *CLV2* gene family reveals *SICLV2* (Solyc04go56640) in a clade with Arabidopsis *AtCLV2* and AON legume *CLV2* genes including pea *PsCLV2* and Medicago *MtCLV2* (Suppl. Fig 1b). An analysis of hydroxyproline O-arabinosyltransferase (*HAPT*) genes revealed tomato *SlFIN* (Solyc11g064850) in a clade with AON legume *RDN1*-like genes from pea *PsRDN1*, Medicago *MtRDN1* and Lotus *LjPLENTY*(Suppl. Fig 1c).

### Pea autoregulation of nodulation mutant shoot and root phenotypes

Since there are no reports of the phenotype of pea *Psnark, Psrdn1* or *Psclv2* mutants in the absence of nodulation, the effects of these mutations on shoot and root growth are confounded by their strong super-nodulation phenotypes. When grown under sterile conditions and no additional nutrients, we found very little difference in shoot or root phenotypes of these mutants compared to wild type (Fig. 1). The only significant findings were a small but significant decrease in tap root length in *Psrdn1* mutants compared to wild type and a small increase in internode length of *Psclv2* mutants compared to wild type. Therefore the large reduction in root and shoot size reported in these mutants in the presence of nodulation (e.g. Foo et al. 2014), appears to be an indirect effect of supernodulation, rather than a direct effect of these genes on shoot or root development.

**Figure 1.**
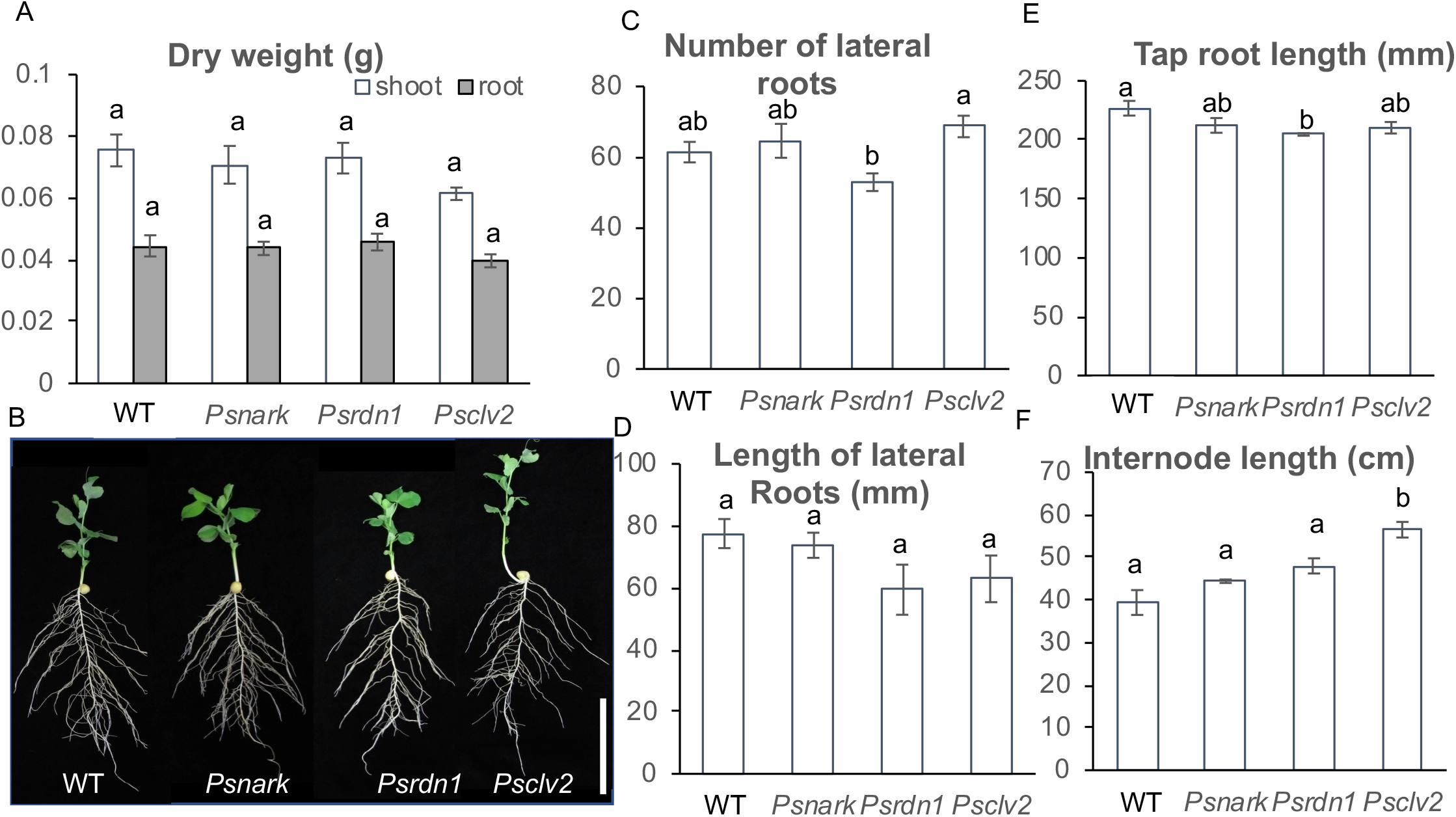
Growth parameters of pea wild type, *Psnark, Psrdn1* and *Psclv2* mutant plants grown in sterile conditions with no additional nutrients for 14 days in pots. A. Dry weight of the shoot and root, B. photo of whole plants (scale bar is 10cm), C. number of lateral roots, D. length of lateral roots, E. tap root length and E. internode length (node 1-5). Data are shown as mean ± SE (n=5-8). Different letters indicate values that are significantly different as assessed by Tukey’s HSD test (P<0.05).

### Mature shoot and root development of *fab, fin* and *clv2-2* tomato mutants

The mature phenotypes of *fab, fin* and *clv2-2* tomato mutants were assessed in pot grown plants (Fig 2). The *fab* mutant displayed a similar shoot and root size to wild type tomato plants. In contrast, the*fin-n2326* and *clv2-2* mutants displayed a significantly smaller root than wild type plants. *clv2-2* mutant plants also displayed a significantly smaller shoot than wild type. This resulted in both *fin-n2326* and *clv2-2* displaying significantly elevated shoot:root ratios compared to wild type plants. Given that in legumes grafting studies have indicated CLV2 acts in the shoot to suppress root nodulation, we also examined shoot and root size in reciprocal grafts between wild type and *clv2-2* plants (Suppl Fig 2). As observed in intact plants, *clv2* self-grafted plants developed smaller shoots and roots than wild type self-grafts. *CLV2* appears to act in a root autonomous manner to control root size, as the small root size was maintained in *clv2* roots grafted to wild type shoots and wild type roots grafted to *clv2* shoots did not display a reduction in root size. Indeed, it is likely that the small shoot size in mature *clv2* plants is due to small root size as *clv2* shoot size was restored by grafting to wild type roots and wild type shoot size was reduced when grafted to *clv2* roots. The root apical meristem size was examined in wild type and various mutant plants grown under similar conditions, although this experiment used an independent allele of *clv2, clv2-5* (Suppl Fig 3). No significant differences in root apical meristem length or area were observed in mutant plants compared to wild type.

**Figure 2.**
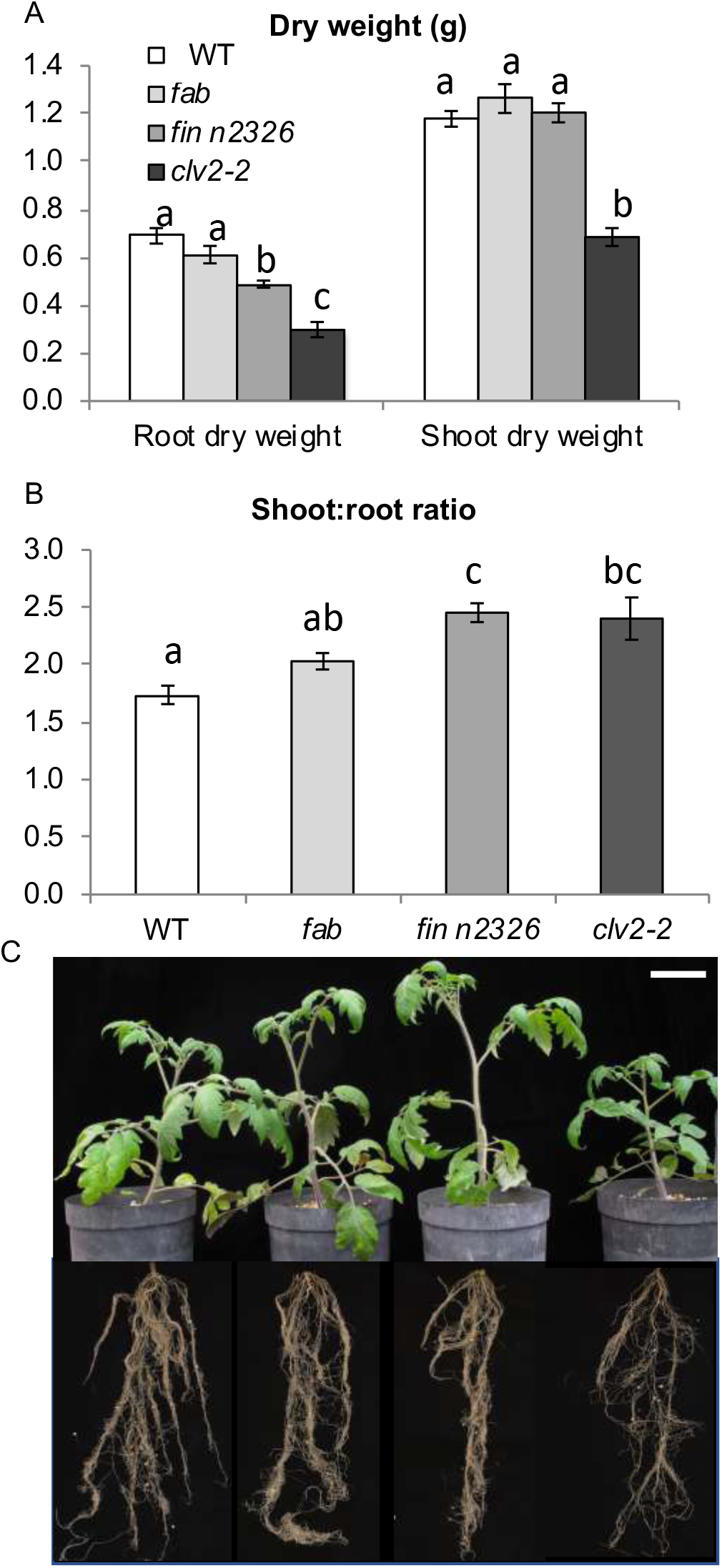
A. Shoot and root dry weights, B. shoot:root ratios and C. photos of 8 weeks old wild type, *fab, fin n2326* and *clv2-2* mutant plants grown in pots. In C. scale bar is 10cm. Data are shown as mean ± SE (n=7-12). Different letters indicate values that are significantly different as assessed by Tukey’s HSD test (P<0.05).

### Shoot and root development of *fab, fin* and *clv2* tomato mutant seedlings under nutrient sufficient conditions

To examine the root development of tomato plants in more detail, tomato seedlings were grown in an in vitro plate system. As observed in mature pot grown plants, there was no significant difference between *fab* mutants and wild type plants in any of the phenotypic characters measured over the 21 days (Fig. 3). In contrast, the *clv2-5* mutant plants had significantly shorter shoots than wild type plants at 10, 14, 17 and 21 days growth. The *clv2-5* mutant seedlings also developed somewhat shorter tap roots early in growth, although this was not significantly different to wild type. Lateral root development in the *clv2-5* mutant plants appeared to be somewhat delayed, with significantly fewer lateral roots after day 10 and 14 and the lateral roots that did develop were significantly shorter than on wild type plants. Plants began to develop tertiary roots after 14 days, and *clv2-5* mutants developed somewhat less tertiary roots although this was not significantly different to wild type plants. Similar trends in tap root length and number of lateral root number were observed in an independent experiment with *clv2-2* mutant plants (data not shown).

**Figure 3.**
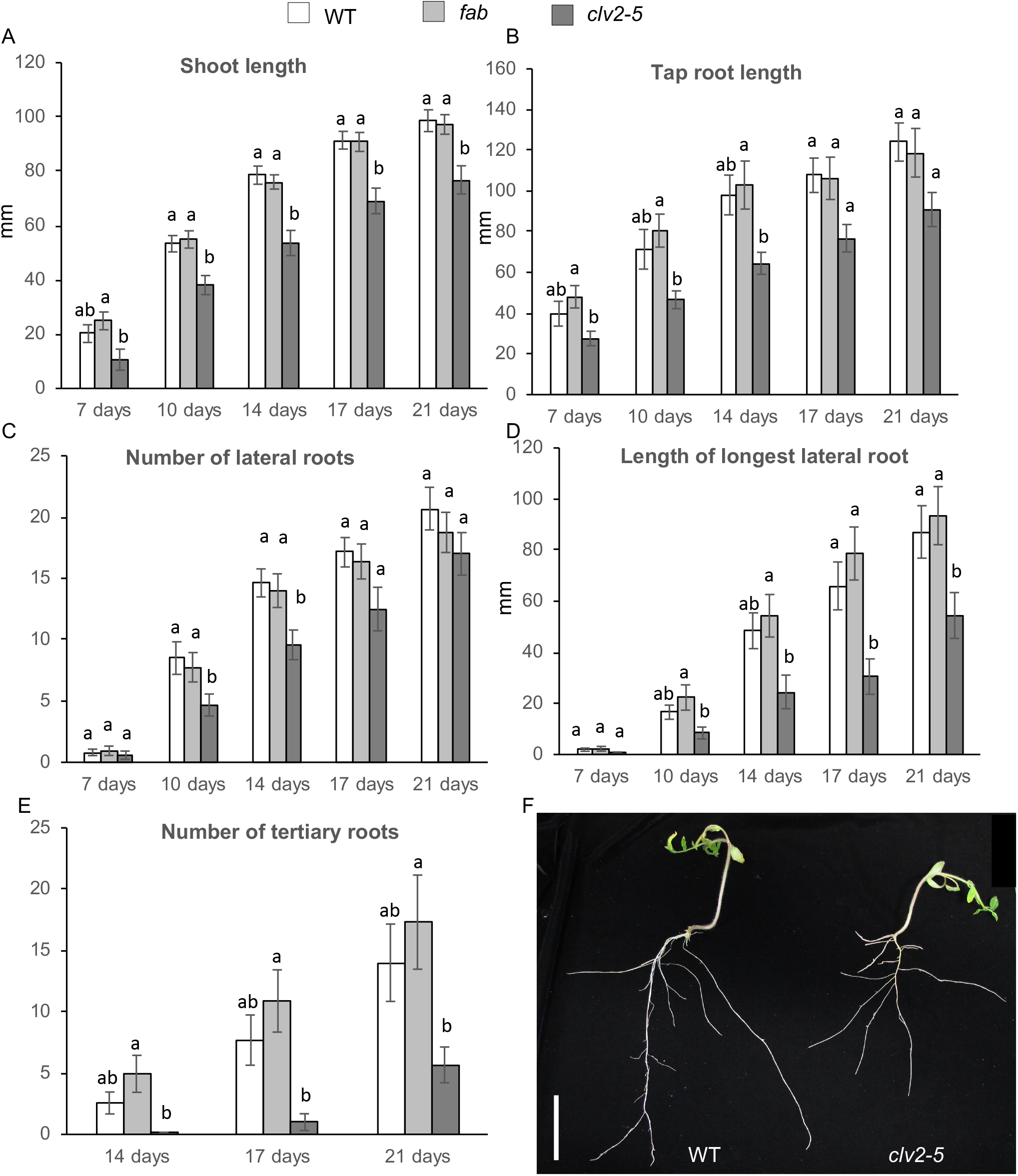
Growth parameters of tomato wild type, *fab* and *clv2-5* tomato mutants growing for 21 days on 2.5 mM P and 5.6 mM N LAN phytagel plates. A. Shoot length, B. tap root length, C. number of lateral roots, D. length of longest lateral root, E. number of tertiary roots and F. photo of 21 days old wild type and *clv2-5* plants, scale bar is 5cm. Data are shown as mean ± SE (n=12-18). Within a time point, different letters indicate values that are significantly different as assessed by Tukey’s HSD test (P<0.05).

Both *fin* mutant alleles (*fin-e4489* and *fin-n2326*) germinated earlier than wild type and initially (days 5-7) had longer tap roots and shoots than wild type (Fig. 4). However, by day 11-14, there was no difference between *fin* mutants and wild type plants in tap root length, shoot length, number of lateral roots or the number of tertiary roots. The only persistent difference between wild type and *fin-n2326* was a reduction in the length of the longest lateral root of *fin n2326* plant*s*, but this difference was not significant in the *fin-e4489* line. In summary, the *fin* mutants germinated earlier than the wild type plants but did not show consistent differences in shoot or root development to the wild type plants.

**Figure 4.**
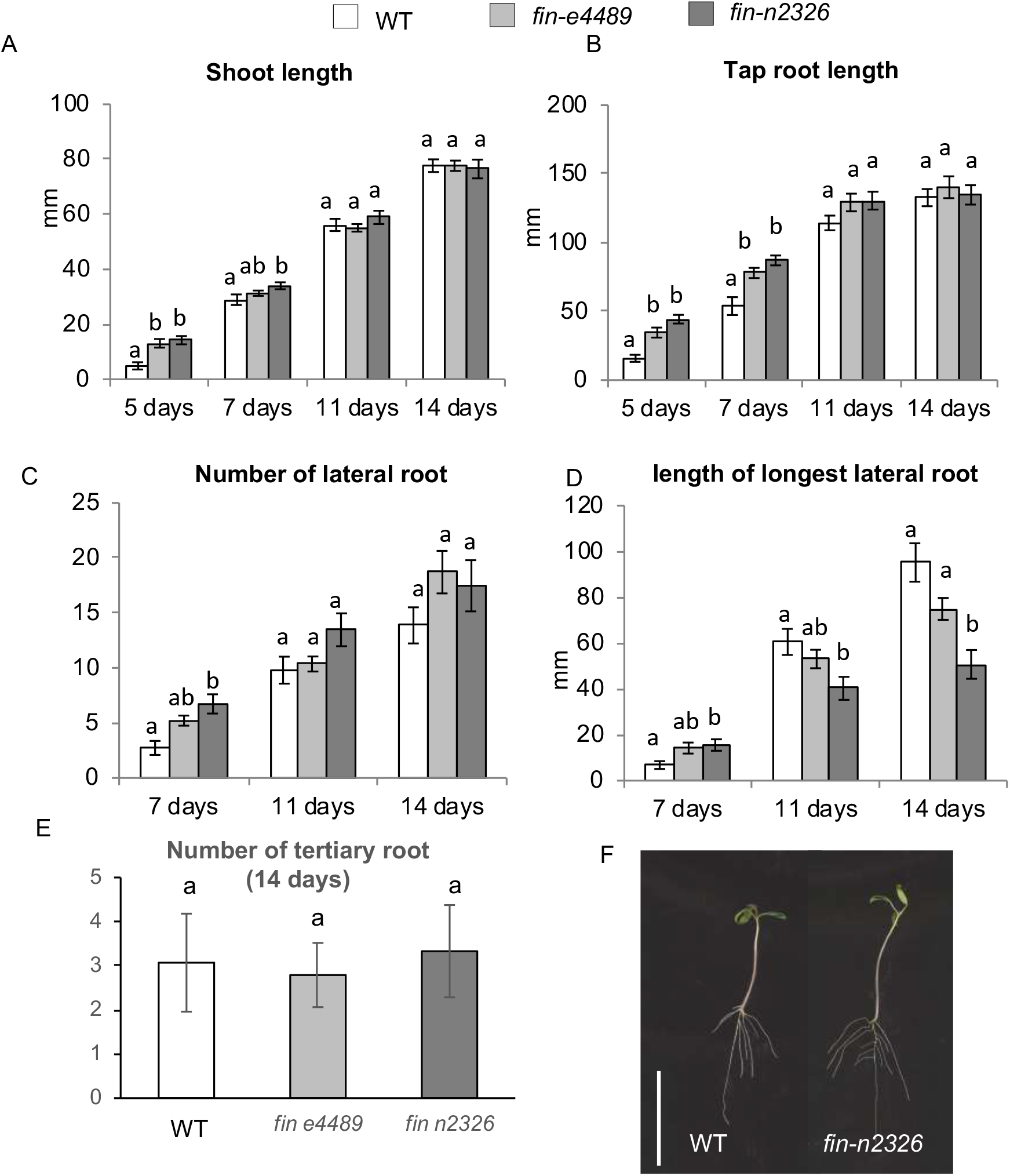
Growth parameters of tomato wild type, *fin-e4489* and *fin-n2326* mutants growing for 14 days on 2.5 mM P and 5.6 mM N LANS phytagel plates. A. Shoot length, B. tap root length, C. number of lateral roots, D. length of longest lateral root, E. number of tertiary roots and F. photo of 14 days old wild type and *fin-n2326* plants. Data are shown as mean ± SE (n=9-13). Within a time point, different letters indicate values that are significantly different as assessed by Tukey’s HSD test (P<0.05).

### Wild type tomato seedlings have altered shoot and root development under low N

To establish the developmental response of tomato roots to nitrogen, we examined wild type tomato plants grown on MGLR media containing a large range of nitrate concentrations (from 10 µM to 7000 µM) as outlined in Araya et al. (2014) (Fig. 5). Significant changes in some aspects of shoot and root development were apparent between N treatments after 10 days of growth. A significant suppression of tap root length by the highest N concentration (7000 µM) was apparent by 10 days (Fig. 5 B). A significant increase in the number of lateral roots was also observed at N concentrations above 100 µM by 10 days, and this was highly significant in N concentrations above 300 µM at 14 days compared to lower N concentrations (Fig 5C). Indeed, at 14 days the number of lateral roots growing under high N conditions (1000 µM and 7000 µM) was nearly double those of the plants growing on 30 and 100 µM N media, and triple those of the plants growing on 10 µM N media. However, no change in the average length of lateral roots was observed in different N treatments (Fig 5.D). These results suggested that low N concentration in the media strongly inhibiting the formation of lateral roots.

**Figure 5.**
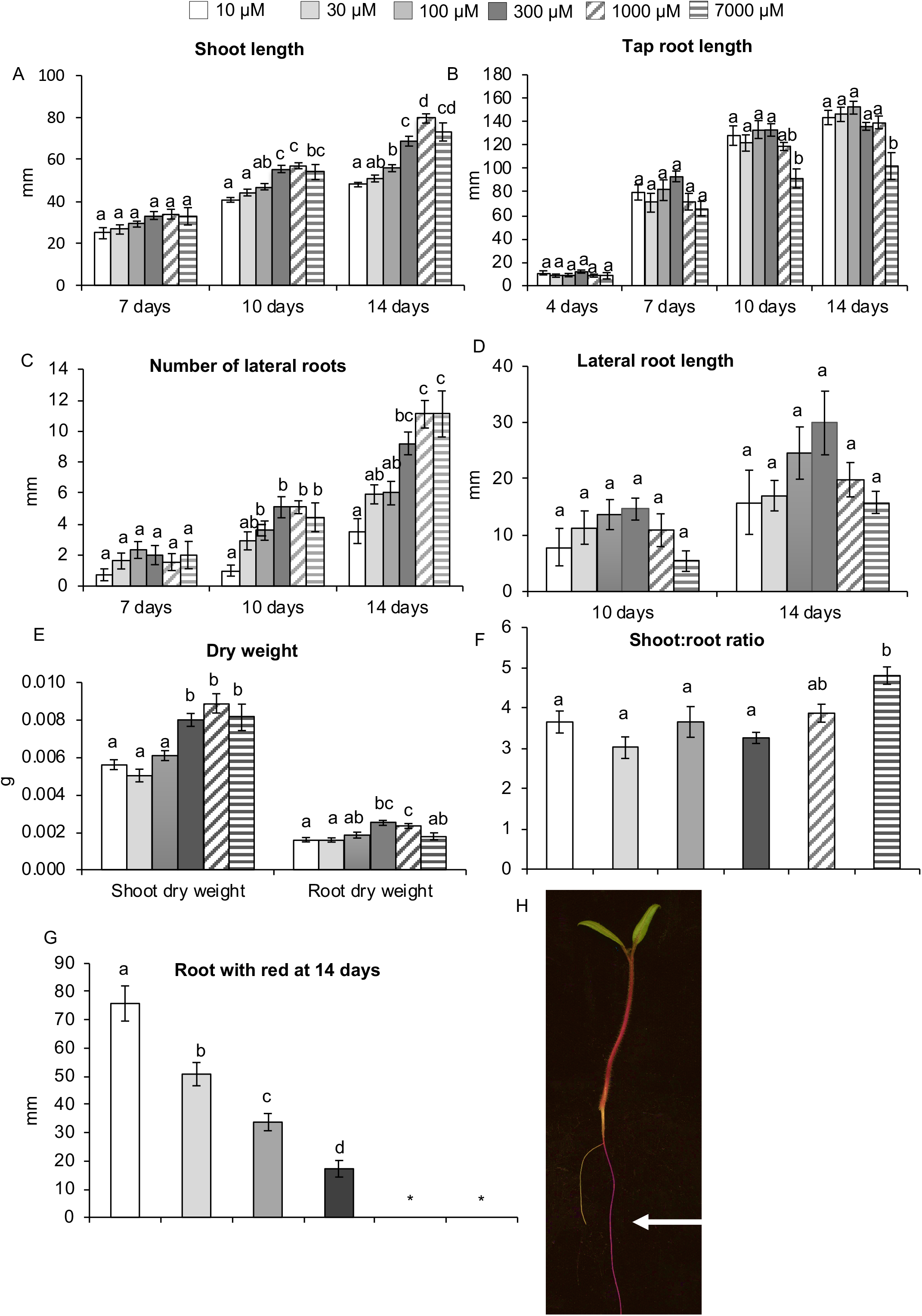
Growth parameters of tomato wild type plants growing for 14 days growing in modified MGRL media with 1.75mM P and different NO_3_^−^ concentrations (ranging from 10 µM to 7000 µM). A. Shoot length, B. tap root length, C. number of lateral roots, D. length of longest lateral root, E. dry shoot and root weight, F. shoot:root ratio, G. length of tap root with red (anthocyanin) colouration (* indicates not detected). Data are shown as mean ± SE (n=15). Within a time point, different letters indicate values that are significantly different as assessed by Tukey’s HSD test (P<0.05).

High N in the media (>300 µM) resulted in longer shoots and higher shoot dry weight than the lower N concentrations (Fig. 5A, E). A small but significant increase in the root dry weight of plants growing in 300 µM and 1000 µM condition was also observed compared to low N conditions. Overall N did not have a dramatic influence on the shoot:root ratio, although plants grown at the highest N (7000 µM) displayed a small but significant increase (Fig. 5F), indicating that plants grown under high N conditions tend to invest more into shoot development. When grown under certain stressful conditions, tomato produces anthocyanin resulting in red colouration of the tissue (Eryılmaz 2006). At 14 days, the length of the tap root with red colouration (anthocyanin) was also measured as indicator of plant stress. The plants growing on the lowest N media (10 µM) had the largest proportion of the tap root displaying red colouration, while the plants grown on N >1000 µM displayed no anthocyanin in the root. In summary, tomato plants grown under low N conditions invested less in shoot development, suppressed lateral root development and accumulated anthocyanins in the root system. In contrast, very high N treatment suppressed tap root elongation and promoted the formation of lateral roots.

### Tomato *fab, clv2* and *fin* mutant seedlings have similar shoot and root development in response to N as wild type

The responses of the *clv2-5, fab* and *fin-n2326* mutants and wild type plants to low, medium and high N concentrations (10µM, 300 µM and 7000 µM) was monitored (Figs 6–8). As observed previously, wild type plants grown under low N displaying suppression of shoot growth and number of lateral roots, promotion in tap root length and accumulation of root anthocyanins. Overall, mutant plants responded to N treatments in a very similar way to wild type as assessed by two-way ANOVA, since across all experiments, there was a significant N treatment effect but there was no treatment by genotype effect. This indicates the mutants are not disrupted in their root morphology N response.

Direct comparison between plants grown on LANS media (2.5mM HPO_4_^-^ and 5600 µM NO_3_^−^; Figs 3–4) and plants grown on MGRL media (1.75mM HPO_4_^-^ and largest N dose 7000 µM NO_3_^−^; Figs 5–8) is challenging due to differences (from approximately 0.5-2-fold) in the concentration of many nutrients. For *fab*, no significant difference from wild type across shoot and root parameters were seen in LANS or MGLR media across all N treatments (no genotype effect as assessed by two-way ANOVA). However, for both *clv2-5* and *fin-n2326* there were subtle differences between phenotypes of these mutants when grown under LANS or MGLR media compared to wild type. By 14 days, *clv2-5* plants grown on LANS media developed shorter shoots, shorter tap roots and fewer lateral roots than wild type (Fig. 3).

**Figure 6.**
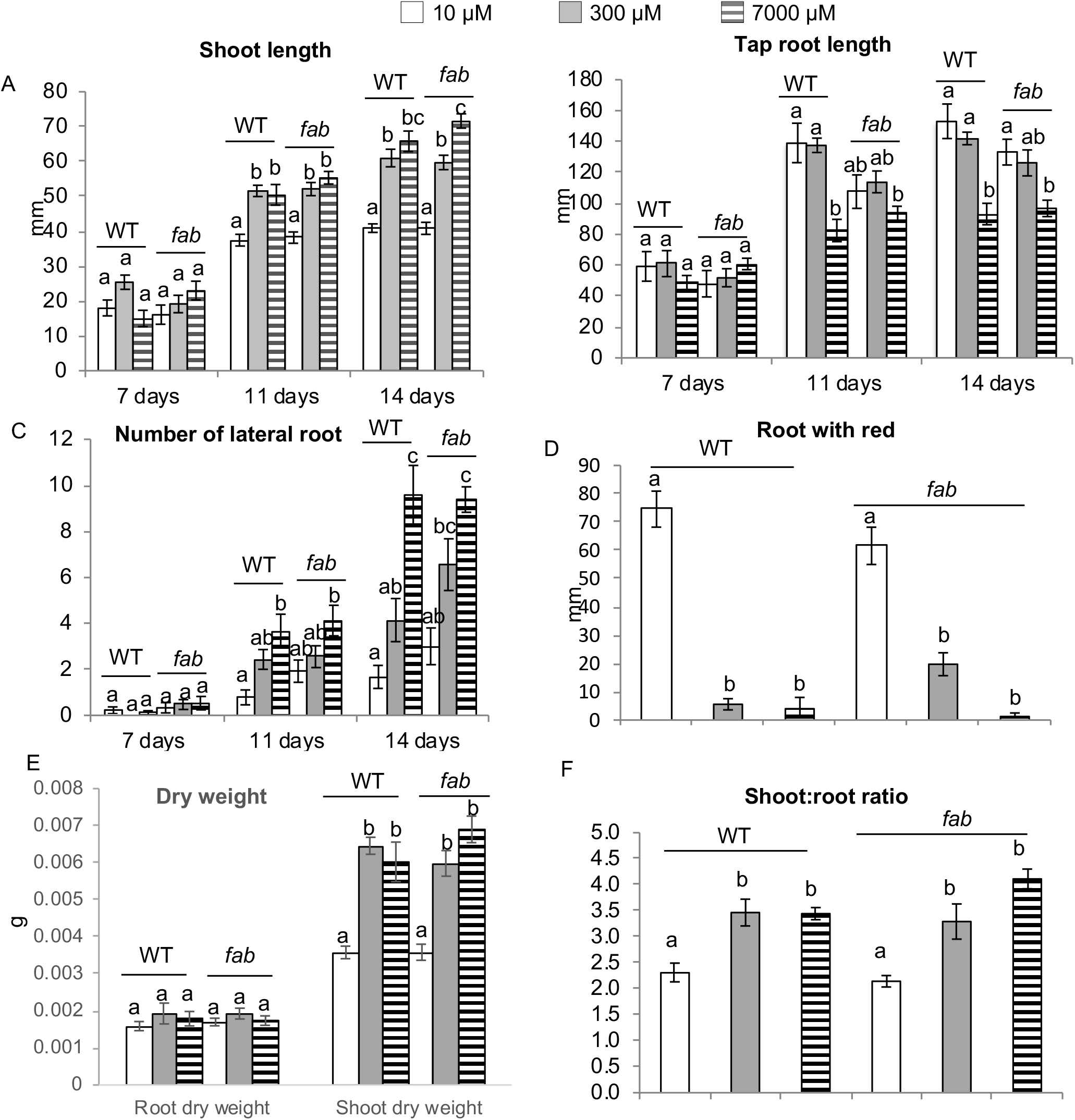
Growth parameters of tomato wild type and *fab* mutant plants growing for 14 days on modified MGRL media with 1.75mM P and different NO_3_^−^ concentrations (10, 300 or 7000 µM). A. Shoot length, B. tap root length, C. number of lateral roots, D. length of tap root with red (anthocyanin) colouration, E. dry shoot and root weight and F. shoot/root ratio. Data are shown as mean ± SE (n=7-17). Within a time point, different letters indicate values that are significantly different as assessed by Tukey’s HSD test (P<0.05).

**Figure 7.**
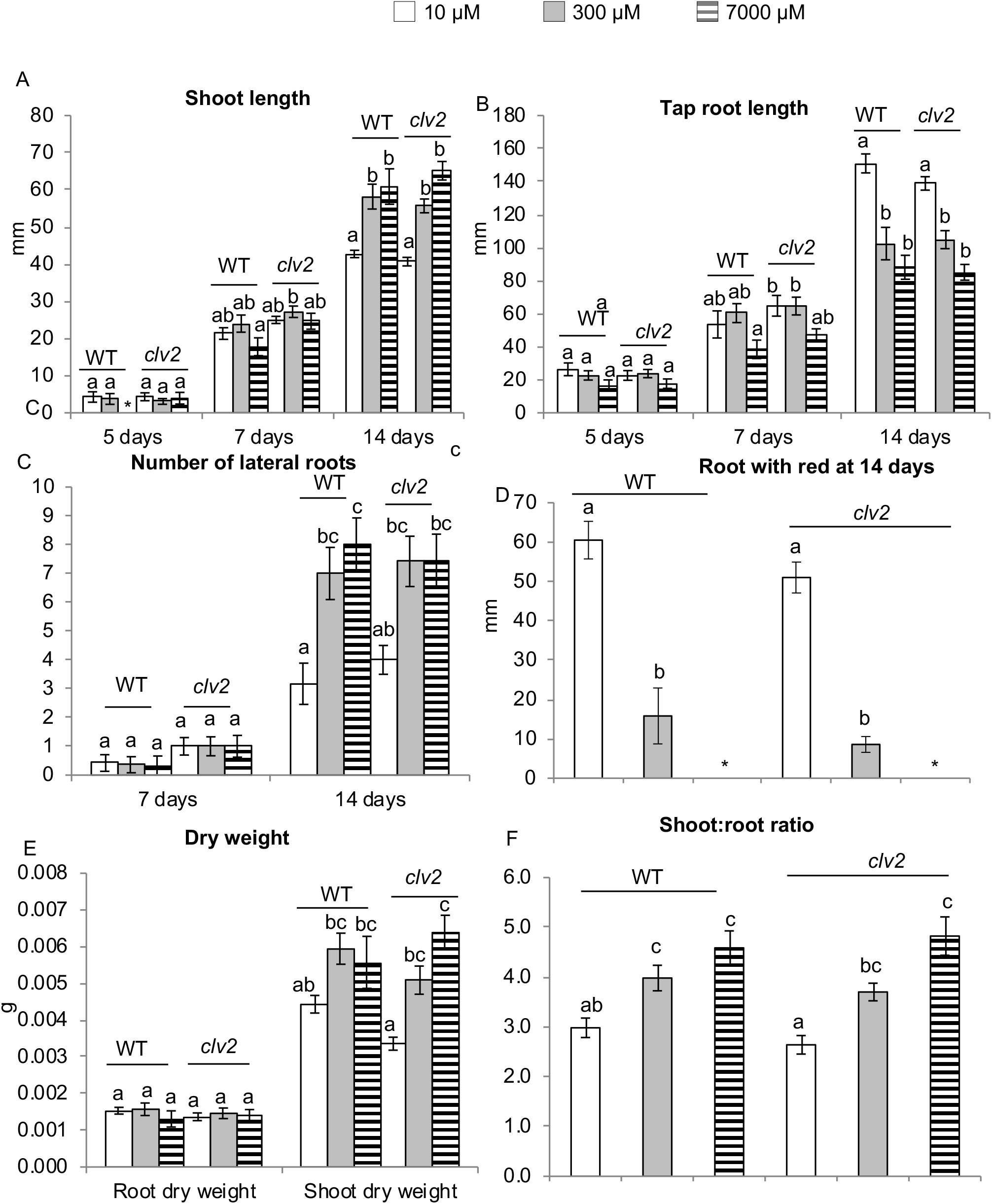
Growth parameters of tomato wild type and *clv2-5* plants growing for 14 days on modified MGRL media with 1.75mM P and different NO_3_^−^ concentrations (10, 300 or 7000 µM). A. Shoot length, B. tap root length, C. number of lateral roots, D. length of tap root with red (anthocyanin) colouration (* indicates not detected), E. dry shoot and root weight and F. shoot:root ratio. Data are shown as mean ± SE (n=9-19). Within a time point, different letters indicate values that are significantly different as assessed by Tukey’s HSD test (P<0.05).

**Figure 8.**
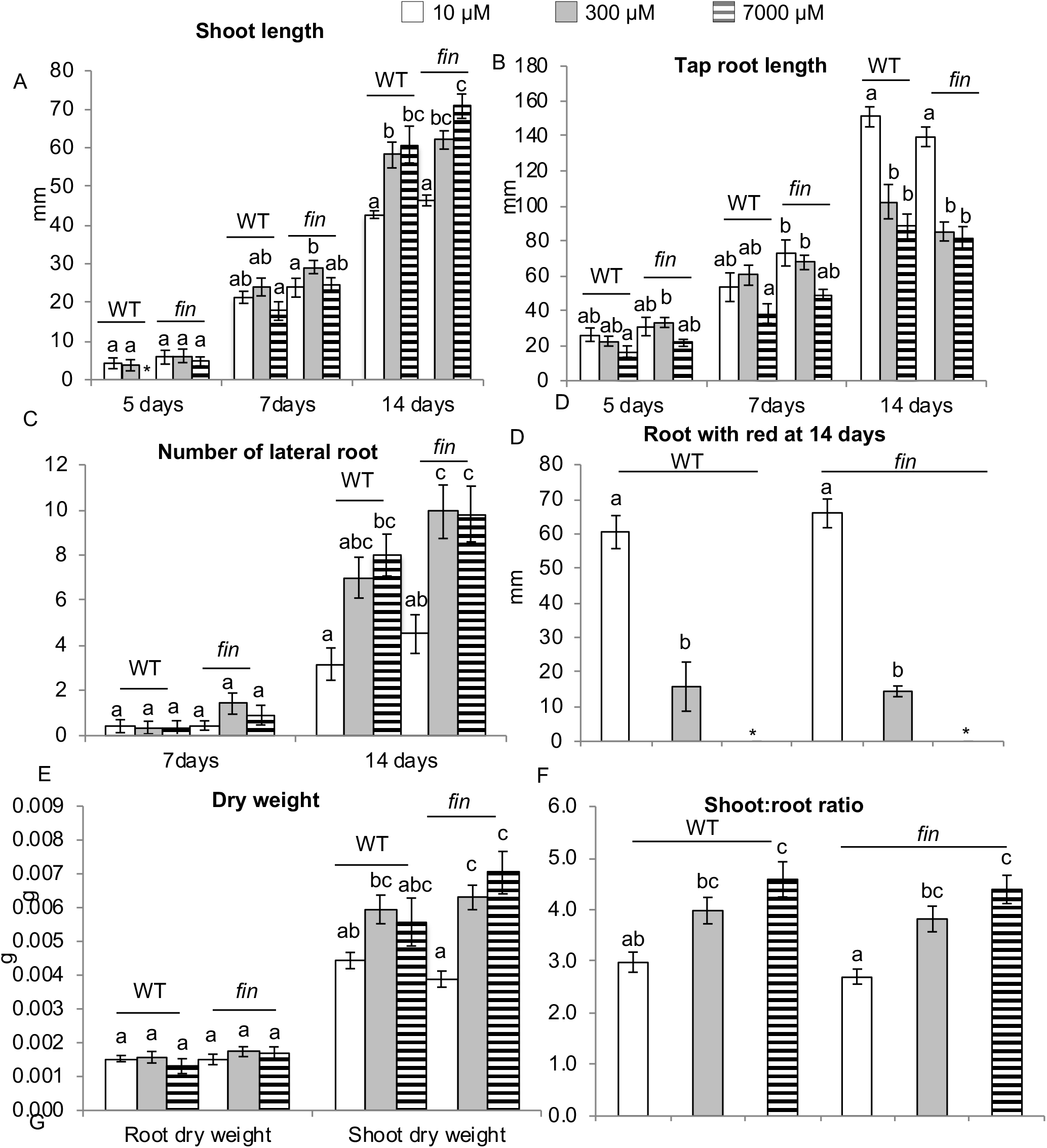
Growth parameters of tomato wild type and *fin-n2326* plants growing for 14 days on modified MGRL media with 1.75mM P and different NO_3_^−^ concentrations (10, 300 or 7000 µM). A. Shoot length, B. tap root length, C. number of lateral roots, D. length of tap root with red (anthocyanin) colouration (* indicates not detected), E. dry shoot and root weight and F. shoot:root ratio. Data are shown as mean ± SE (n=9-19). Within a time point, different letters indicate values that are significantly different as assessed by Tukey’s HSD test (P<0.05).

However, in the experiment performed on MGLR media no significant change in these parameters were observed between *clv2-5* and wild type plants at any N treatment (no genotype effect as assessed by two-way ANOVA). For *fin-n2326*, similar trends but subtle differences were observed between LANS and MGLR experiments. When grown in MGLR media, *fin-n2326* mutant plants showed small but significant increases in tap root length and shoot length (5, 7, 14 days) and the number of lateral roots at 14 days compared to the wild type plants (significant genotype effect as assessed by two-way AONVA), although these differences were small (<10%) and would only have a minor impact on plant development overall. In contrast, in plants grown in LANS *fin-n2326* plants only displayed longer tap roots and shoots than wild type from 5-7 days and no significant change in the number of lateral roots. These minor differences illustrate that the subtle changes in root development sometimes observed in *clv2* and *fin* mutants seedlings are clearly contingent on specific growth conditions.

## Discussion

In this paper, we expanded our understanding of the role of CLV1 in root response to N by examining the role of CLV1, CLV2 and HPAT homologs in pea and tomato. Although tomato mutants disrupted in *Sl*CLV2 and HPAT *Sl*FIN displayed some modification of root architecture, suggesting a role for these proteins in root development of tomato, we found no evidence for these proteins or CLV1 orthologue *Sl*FAB in the control of N regulation of tomato root development. No consistent effect on root architecture of seedlings was observed in homologous pea mutants grown in the absence of external nutrient. Thus, the role of these genes in specific aspects of root development may be relatively species specific and CLV1 does not appear to be a central regulator of N response of roots across species.

In pea, disruption of genes important for the autoregulation of nodulation pathway (*PsNARK, PsCLV2* and *PsRDN1*) did not result in any major change in root development in the absence of nodulation and external nitrogen supply. Thus, the substantial reduction in root size previously reported in these pea mutants (e.g. Foo et al. 2014) appears to be largely an indirect effect of the supernodulation phenotype that develops in the presence of rhizobia. The small but significant increase in internode length observed in *Psclv2* mutant lines (Fig. 1) is consistent with the influence of this gene on shoot development, in particular the development of a fasciated shoot apex in *Psclv2* mutants (Krusell et al. 2011) and in homologous mutants in other legumes and non-legumes (e.g. Krusell et al. 2011; Xu et al. 2015). Mutants orthologous to *Psnark* in other legume species (*Mtsunn/Ljhar1/Gmnark*) have been reported to display changes in root development under uninoculated conditions, so this lack of phenotype in pea mutants (Fig. 1) is intriguing. It is interesting to note that although *CLV1-like* and *RDN1-like* genes in the non-legumes Arabidopsis, rice, tomato have clear roles in shoot apical meristem development (e.g. Clark et al. 1997; Suzaki et al. 2004; Xu et al. 2015) there is no phenotypic evidence that these genes play similar roles in legumes (e.g. Sagan 1996; Schnabel et al. 2011; Searle et al. 2003). Therefore, the specific function of these genes in shoot and root development may vary between species.

A species specific role for *CLV1-like, CLV2* and *RDN1-like* genes in root development is also supported by the experiments conducted with tomato mutants. We found no evidence for a role for *FAB* in tomato root development either at seedling or more mature plant stage. Indeed, although tomato seedlings responded strongly to differences in N conditions, we also found no evidence that *FAB* plays a role in the N regulation of root architecture. This is in contrast to similar studies conducted with Arabidopsis *clv1* and *M. truncutula sunn* mutants which indicated these genes play a key role in root development including the response to N, indicating that the role of CLV1-like genes in roots may be species specific. The role for *RDN1-like* genes in root development has received less attention, but there is some evidence that at least in *M.truncutula rdn1* mutants may have altered root development and altered response to N (Goh et al. 2018). We did find that mature tomato *fin* mutants developed somewhat smaller root systems with no corresponding change in shoot size, indicating a change in shoot:root partitioning. However, we detected only small changes (<10% change compared to wild type) in *fin* seedling root architecture under some growth conditions and like FAB, we found no evidence for a role for FIN in the N regulation of root development. Thus, although *rdn1-like* mutants in both *M. truncatula* (Goh et al. 2018), *L. japonicus* (Yoshida et al. 2010) and tomato developed a smaller root system the role of this gene in the N response did not appear to be conserved.

Of all the mutants assessed, the root development of tomato *clv2* mutants was the most significantly affected. Mature pot grown *clv2* tomato mutant plants developed significantly smaller roots and shoots than wild type plants. Grafting studies indicated *CLV2* appears to act in the root to control root size, and the smaller shoot size of *clv2* mutants may be an indirect effect of this reduced root size. The fewer lateral roots observed in *clv2* mutant seedlings that were cultivated on LANS media, suggested this may be a mechanism through which *CLV2* positively influences root size. In some studies conducted with homologous mutants in other species, differences in specific cell layers of the root apical meristem have been observed (e.g. Stahl et al. 2013; Whitewoods et al. 2018). While a detailed examination of these phenotypes was beyond the scope of this study, an examination of root apical length and area did not indicate any significant differences between the tomato *fab, clv2* and *fin* mutants and the wild type plants (Suppl Fig 3). In many species, significant insights into the role of CLE peptide signalling in root development has been enabled by peptide application studies and/or *cle* mutant or transgenic studies and this could be an approach for future studies.

## Abbreviations

CLV1: CLAVATA1
CLV2: CLAVATA 2
HPAT: hydroxyproline O-arabinosyltransferase
N: nitrogen

## Author contributions

EF, CW and JBR conceived of the project. CW performed experiments and analysed data. CW, EF and JBR prepared the manuscript.

## Acknowledgements

We thank Prof Zachary Lippman and Dr Choon-Tak Kwon for the kind gift of the *fab, clv2* and *fin* mutant lines and for helpful discussions. We thank Fu Rong Mah, Michelle Lang, Tracey Winterbottom and Valerie Hecht (University of Tasmania) for technical assistance.

## Supplementary Figure legends

Suppl Figure 1. Phylogenetic tree of (A) CLV1-like, (B) CLV2, and (C) HPAT family proteins. The sequences included are from tomato (*Solanum lycopersicum*, Slj, *Pisum sativum* (Ps), *Lotus japonicus* (Lj), *Medicago truncatula* (Mt), soybean (*Glycine max*, Gm), *Oryza sativa* (Os) and *Arabidopsis thaliana* (At). Bootstrap values are indicated.

Suppl Figure 2. Shoot and root fresh weight in reciprocal grafts between *clv2-2* and wild type tomato plants 6 weeks after grafting. Data are shown as mean ± SE (n=14 plants per graft). Within a parameter, different letters indicate values that are significantly different as assessed by Tukey’s HSD test (P<0.05).

Suppl Figure 3. Root apical meristem measurements (RAM) of 4 weeks old wild type, *fab, fin-n2326* and *clv2-5* mutant tomato plants grown in pots. A. RAM length, B. RAM area and C. photo of stained root tip. Data are shown as mean ± SE (n=3 plants).

## Data availability statement

The data that support the findings of this study are available from the corresponding author upon reasonable request.

